# Activation of the *Yersinia* type III secretion system induces large-scale chromosomal and virulence plasmid DNA rearrangements

**DOI:** 10.1101/2025.02.04.636459

**Authors:** Francesca Ermoli, Christoph Spahn, Ismath Sadhir, Helge B Bode, Andreas Diepold

**Author notes:** Corresponding author: Andreas Diepold.

## Abstract

The type III secretion system (T3SS) is used by Gram-negative bacteria, including important pathogens, to manipulate eukaryotic target cells by injecting effector proteins. Type III secretion and bacterial physiology are known to be tightly interconnected and influence each other. Most notably, secreting cells undergo growth arrest in the T3SS model organisms *Yersinia*, *Salmonella* and *Shigella*. The molecular basis of this phenotype, referred to as secretion-associated growth inhibition, is debated. In *Yersinia*, T3SS genes are encoded extra-chromosomally in the plasmid of *Yersinia* virulence (pYV), whose copy number increases upon induction of T3SS secretion. In this study, we characterize the link between T3SS activity and subcellular organization by localizing and quantifying the pYV and chromosomal DNA in *Yersinia enterocolitica*. We find that activation of secretion not only increases the number of pYV plasmids per bacterium, but that the plasmids also move towards the membrane and poles. This relocalization is not caused by transertion (coupled transcription, translation and translocation) of effectors, but part of a broader DNA rearrangement, leading to a distinct relocalization of chromosomal DNA to mid-cell. We hypothesize that these striking DNA rearrangements occurring during secretion are a main factor in the secretion-associated growth inhibition of pathogenic bacteria.

## Introduction

The type III secretion system (T3SS) is used by many Gram-negative bacteria to manipulate the host during the establishment of an infection [1–4]. The T3SS, also called injectisome, is a highly conserved multi-protein double membrane-spanning nanomachine that enables the direct injection of effector proteins from the bacterial cytosol into the target host cell [5]. While the injectisome is conserved in different bacteria, its export substrates, the effector proteins, greatly vary among species and can exert a variety of functions supporting the pathogen-specific infection strategy [5–7]. In the three *Yersinia* human pathogenic species *Y. pestis*, *Y. enterocolitica,* and *Y. pseudotuberculosis,* the 70kb <u>p</u>lasmid for *<u>Y</u>ersinia* <u>v</u>irulence (pYV) [8] encodes for the T3SS machinery, the effectors, called *<u>Y</u>ersinia* <u>o</u>uter <u>p</u>roteins (Yops), and T3SS chaperones, which are required for the export of some effectors. The *Yersinia* T3SS is essential for evading the immune system by preventing phagocytosis and the activation of the inflammatory cascade [9]. The two main stimuli regulating T3SS secretion are temperature and host-cell contact. Expression and assembly of the T3SS machinery occurs upon temperature shift to the host temperature (37°C) via a complex regulatory cascade ultimately inducing the synthesis of the main transcriptional activator of the system, VirF/LcrF (reviewed in [10]). T3SS secretion is consequently activated by contact with the target host cell membrane, a condition efficiently mimicked *in vitro* by Ca^2+^ depletion in the culture medium [11].

Activation of secretion leads to a strong upregulation of the production of effectors [12–15], but also a roughly two-fold upregulation of the levels of injectisome components and as a result twice as many injectisomes per bacterium [16]. This is likely caused by a similarly increased number of pYV plasmids. In *Y. pseudotuberculosis*, the pYV plasmid copy number (PCN) increases from about one chromosome equivalent to 1.5-2 upon host entry (37°C) and further to 3-4 over time [17,18]. The PCN is modulated by the plasmid replication initiator RepA via the CopAB system. CopB inhibits *repA* transcription by repressing its promoter, while CopA is an antisense RNA preventing *repA* translation by interacting with mRNA ribosome binding site (RBS) [19,20]. Under T3SS-inducing conditions, increased pYV PCN is triggered by CopAB downregulation [18].

Besides the copy number, the localization of DNA can have important consequences. A recent study demonstrated that the expression and assembly of the T3SS2 of *Vibrio parahaemolyticus*, which is encoded on the chromosome, is likely mediated via transertion [21,22]. Transertion generates a dynamic link between the genomic locus and the inner membrane via coupling of transcription, translation and insertion (or translocation) of proteins into (or across) the inner membrane. Particularly for low-abundant structures, such as the *V. parahaemolyticus* T3SS2, the spatial proximity of the gene and transcript to the site of insertion/translocation can enhance efficiency and reduce the metabolic burden. Disruption of transcription or translation leads to a rapid contraction of the bacterial nucleoid and loss of the putative link to the membrane [22,23]. While evidence for the existence of transertion is growing, its role and effect on cell physiology is still unclear.

T3SS secretion can distinctly alter bacterial physiology. One of the most drastic effects is that T3SS-active cells display a severe growth retardation. This phenomenon, termed secretion-associated growth inhibition (SAGI), was the phenotype that initially led to the discovery of the T3SS, when researchers noted a strong growth inhibition specific to virulent *Yersinia* (now known to be equivalent to the presence of the T3SS) at 37°C (T3SS expression) in the absence of Ca^2+^ (T3SS activation) [24–29]. Besides *Yersinia*, SAGI has been shown to occur in two other main T3SS model organisms, *Salmonella enterica* and *Shigella flexneri* [30–32]. However, it is still unclear if SAGI is a mere consequence of the energy expenditure for synthesis and export of the T3SS machinery and effectors, or whether it is an actively regulated phenotype, as suggested by the fact that the levels of key players of chromosome replication and segregation are unchanged in secreting cells [33] and that growth and division are reinitiated very quickly, once secretion is stopped [34].

To investigate the effect of T3SS activation on bacterial cell biology, we compared key cellular parameters in secreting and non-secreting *Y. enterocolitica*. Strikingly, live-cell and super-resolution microscopy revealed a drastic reorganization of the bacterial DNA: Activation of secretion led to a strong condensation of the chromosomal DNA at mid-cell within 30 to 60 min, whereas the virulence plasmid not only increased in copy number but relocalized towards the bacterial membranes. While effector mRNA was strongly upregulated under secreting conditions, it was not enriched at the bacterial membrane, arguing against the transertion of effectors. This profound reorganization of bacterial DNA upon activation of secretion could be a missing link in the secretion-associated growth inhibition of secreting bacteria.

## Results

### pYV copy number is affected by temperature and T3SS activation

To investigate the influence of activation of the type III secretion system on pYV plasmid copy-number (PCN), we quantified the PCN at 28°C and 37°C (host temperature) in non-secreting and secreting conditions. Secretion can be controlled by the Calcium concentration of the medium in *Yersinia* [35]. However, to exclude any unspecific effect of the medium, we used a T3SS-deficient (Δ*sctD*) and a constitutively T3SS-active (Δ*sctW*) strain in this study. SctD is an essential inner-membrane structural component of the T3SS, while SctW is the main component of the gatekeeper complex, responsible for T3SS secretion inhibition prior to host-cell contact [11,36]. From now on, we will refer to these strains as non-secreting and secreting, unless differently specified.

Previous studies have shown that in *Y. pseudotuberculosis*, the copy number of the pYV virulence plasmid increases upon host entry (temperature shift to 37°C) and further upon activation of T3SS secretion [37,38]. To test if this is the case in *Y. enterocolitica* as well, we measured the ratio of pYV copy number and chromosomal copy number by qPCR, using the pYV-encoded T3SS transcription factor *virF*/*lcrF* and the chromosomally encoded *gyrB* genes as templates. Similar to the results in *Y. pseudotuberculosis* [17], pYV PCN (more precisely, the *virF*/*gyrB* ratio) increased from around 1 (28°C) to 2 at the host temperature (37°C, non-secreting) and further to 3-4 upon activation of secretion (37°C, secreting) (**Fig. 1A**). To visualize the cellular distribution of pYV plasmids and to evaluate the heterogeneity of the PCN in the different conditions on a single-cell level, we inserted a *parS_P1_* sequence into a region of the pYV plasmid that is not required for T3SS function (the T3SS effector YopE) (*yopE_1-53_*-*parS_P1_*). Expression of an mScarlet-I fusion to the corresponding ParB protein, mScarlet-I-ParB_P1_, from an inducible plasmid (**Suppl. Fig. 1A**), led to the detection of fluorescent foci. We chose this approach over using the native ParB/*parS* system to not interfere with pYV replication and segregation. Functionality and stability of the mutants and specificity of the mScarlet-I-ParB_P1_-*parS_P1_* spot formation was confirmed (**Suppl. Fig. 1BC**). Single-cell quantification of parB_P1_ foci yielded results highly comparable with the qPCR data (**Suppl. Fig. 2**). It confirmed the effect of both temperature and T3SS activation on the PCN and highlighted a considerable heterogeneity in pYV copy number (**Fig. 1B**).

**Figure 1 –.**
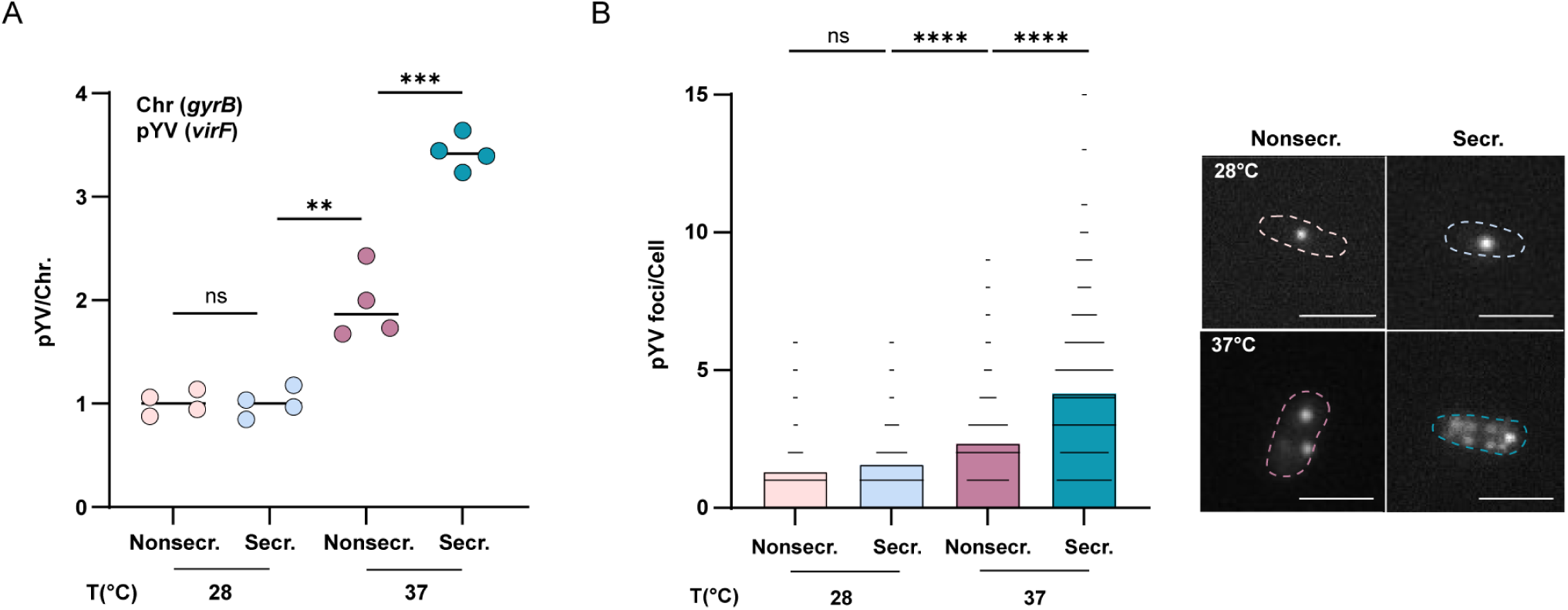
pYV plasmid copy number is influenced by temperature and activity of the T3SS. **A)** Plasmid copy number expressed as pYV (*virF*) / number of chromosome equivalents (*gyrB*) in non-secreting (nonsecr., Δ*sctD*) and secreting (secr., Δ*sctW*) bacteria at 28°C and 37°C, as determined by qPCR. Statistical analysis was performed using a non-parametric t-test, ns, *p>*0.05; **, *p*<0.01; ***, *p*<0.001. *n*=4, circles indicate individual biological replicates, black bars denote the mean. **B)** Quantification of pYV (*yopE_1-53_-parS_P1_*) spots per bacterium, visualized by fluorescence microscopy in strains expressing mScarlet-ParB_P1_ at different conditions as in A). Sample micrographs depicted on the right, scale bar, 2 µm. Statistical analysis was performed using a non-parametric t-test, ns, *p>*0.05; ****, *p*<0.0001, black spots indicate individual measurements. *n*=3.

### T3SS activation induces a re-localization of the pYV plasmid towards the membrane

Next, we assessed the localization of the pYV plasmid in secreting and non-secreting bacteria. First, we determined the localization along the width of the cell by measuring the distance of the plasmid from the longitudinal centerline. The distribution of the T3SS component SctQ (EGFP-SctQ) [39] was taken as a control for membrane localization. In T3SS-active cells, the pYV plasmids localized significantly closer to the membrane than in non-secreting cells (**Fig. 2A**). As spot distributions were symmetric, statistics were performed on the absolute values. Different from the PCN, which increased upon both temperature shift and T3SS triggering (**Fig. 1**), the pYV plasmid relocalization was only observed upon induction of secretion (**Fig. 2A**). To test whether the observed pYV reorganization upon T3SS activation could be explained as a mere consequence of PCN increase, we determined the pYV distribution in cells displaying one or two spots in the previously tested conditions. In both groups, the pYV plasmids of T3SS-active cells were significantly shifted from the centerline to the membrane compared to the other conditions tested (**Fig. 2B**).

**Figure 2 –.**
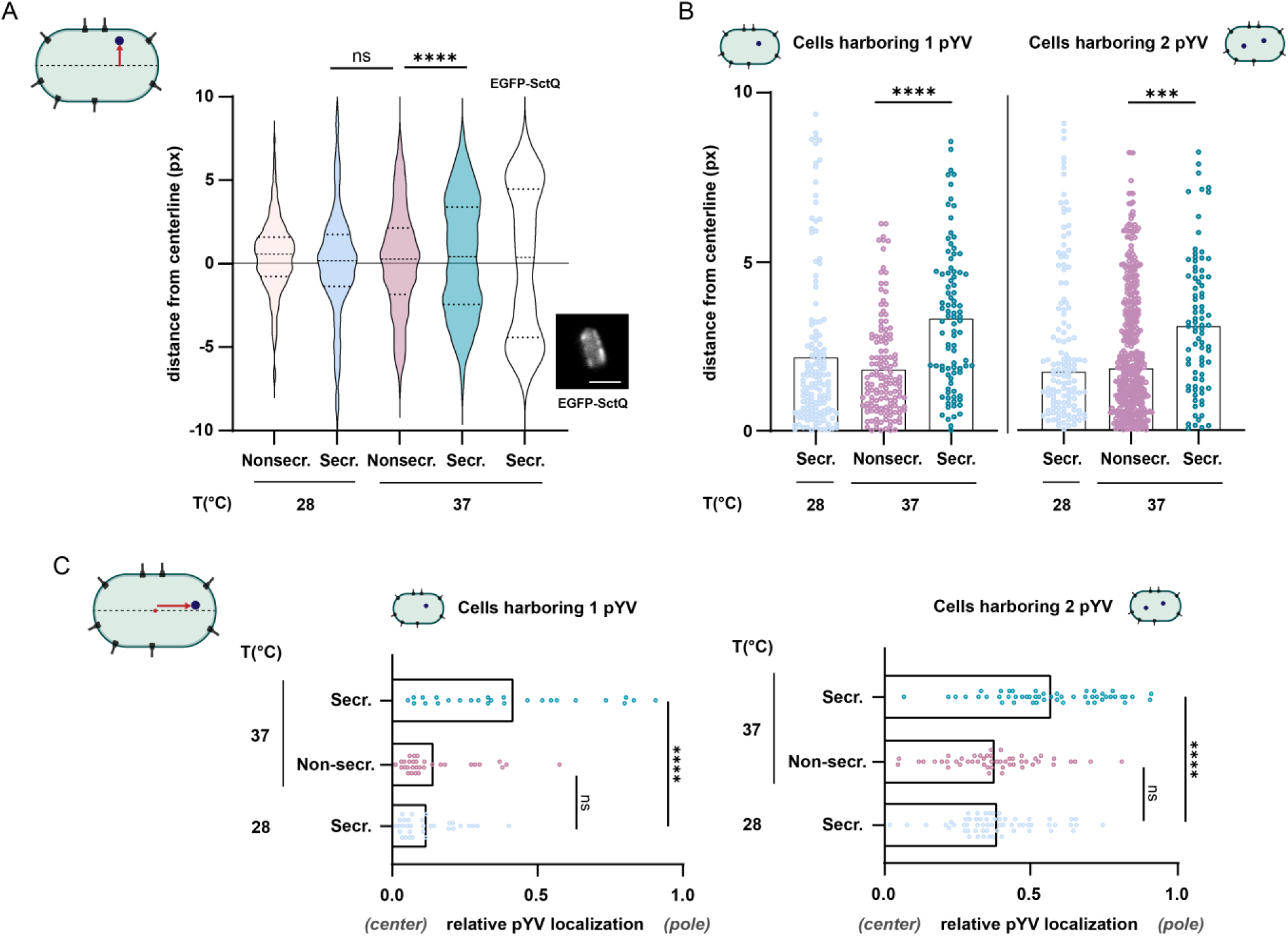
The *Yersinia* virulence plasmid relocates towards the membrane and poles in secreting bacteria. **A)** Single-cell pYV plasmid localization (*yopE_1-53_-parS*) measured as distance from the centerline (px) in the indicated conditions. Statistical analysis was performed on absolute values using a non-parametric t-test, ns, *p*>0.05; ****, *p*<0.0001, dotted lines represent median and quartiles. Representative fluorescence micrograph showing EGFP-SctQ as a control for membrane localization. **B)** Single-cell pYV plasmid localization as above for cells displaying one spot (left) or two spots (right). Statistical analysis was performed using a non-parametric t-test, ***, *p*<0.001; ****, *p*<0.0001; spots indicate single measurements. *n*=3 for all panels. **C)** Single-cell pYV plasmid localization along the longitudinal axis for cells displaying one (left) or two spots (right). To account for differences in cell length, the relative localization of plasmids is displayed, 0 = center of cell, 1 = cell pole.

Next, we quantified the localization of pYV plasmid along the longitudinal axis (central vs. polar localization). As secretion influences the length of bacteria, we determined the relative localization of pYV plasmids on the center to pole line (0 = central localization, 1 = polar localization). In secreting bacteria, the virulence plasmids were strongly shifted from the center towards the poles of the bacteria. Like in the previous experiment, we also compared cells with one or two foci in the different conditions and found that the relocalization of pYV plasmids towards the poles was independent of the increase in plasmid number (**Fig. 2C**).

### Synthesis and secretion of the effector YopM are uncoupled

It has been speculated that T3SS-harbouring bacteria may benefit from using transertion during injectisome assembly [22] or substrate secretion. Transertion couples co-transcriptional translation and protein insertion at the site of action or export to optimize synthesis efficiency [40–42]. As this process requires the presence of the target gene at the site of protein insertion, we wondered whether the observed relocalization of pYV plasmids under secreting conditions is caused by transertion (**Fig. 2**). As transertion requires mRNA to be localized near the membrane, we investigated the localization of the mRNA of the T3SS effector YopM by RNA fluorescent *in situ* hybridization (RNA-FISH). This candidate is particularly suited because it does not harbor a cognate chaperone [43,44]. To ensure single-particle detection and allow for the co-localization of the transcript with bacterial DNA and membrane, we used single-molecule super-resolution microscopy (SMLM). To visualize transcripts, we used DNA points accumulation for imaging in nanoscale topography (DNA-PAINT) coupled with RNA-FISH (super-resolution single-molecule RNA FISH, sr-smRNA-FISH) (**Fig. 3A**). In this method, the transient hybridization of a target complementary strand (docking strand) and a short dye-labeled oligonucleotide (imager strand) enable multiplexing and high accuracy during super-resolution imaging [45]. The localization precision was 10 - 11 nm throughout all measurements as determined by Nearest Neighbor Analysis (NeNA) [46], translating to a spatial resolution of ~ 25 nm. As positive control of the system, the eubacteria-specific rRNA probe EUB338 was used, while a sample processed without any mRNA targeting probe was used as negative control (**Suppl. Fig. 3**). DNA and membranes were visualized using conventional PAINT, using JF_646_-Hoechst and Potomac Gold as transiently binding probes [47]. All three super-resolved channels were overlaid to visualize the spatial context of *yopM* mRNA, the nucleoid and the cell envelop and to assess mRNA enrichments in specific cellular compartments (**Fig. 3B**). *yopM* mRNA abundance strongly increased in secreting conditions, with transcripts being excluded from the nucleoid area (**Fig. 3B**, yellow). However, we did not observe a specific enrichment at the cell membrane, but an apparently higher density at the cell poles and between the sister chromosomes (**Fig. 3B, Suppl. Fig. 4**). This argues against transertion as a strategy used for YopM synthesis and secretion. The high resolution of the sr-smRNA-FISH measurements allows for quantification of transcript abundance. We performed cluster analysis using Density-Based Spatial Clustering of Applications with Noise (DBSCAN) [48] with experimentally derived parameters (**Suppl. Fig. 5**) and subsequent filtering (see Methods) to extract the number of clusters per cell as a proxy for mRNA copy number. Here, the number of transcripts under secreting conditions correlated with cell size (**Fig. 3C**). Under non-secreting conditions, only slightly more transcripts were detected than in the negative control (2.4 ± 2.8 vs 1.2 ± 1.1 clusters, s.d., n = 176 and 33 cells for non-secreting cells and the control). 90 min after induction of secretion, we observed a ~13-fold increase in transcript abundance, with a high variability between single cells (31 ± 13 clusters, s.d., n = 531 cells from 4 replicates) (**Fig. 3D**), while the mean values of individual replicates were highly reproducible (31 ± 4 clusters, s.d.). Similar trends were seen for the cluster density (# clusters per µm^2^ cell area) (**Suppl. Fig. 6**). At the sub-cellular level, the cluster density under secreting conditions appears to be higher at the poles, where we also observed a higher abundance of ribosomes, in accordance with previous results [49] (**Fig. 3E**). We thus conclude that the *yopM* mRNA freely diffuses in the cytosol and gets captured at ribosome-dense regions by uncoupled translation.

**Figure 3 –.**
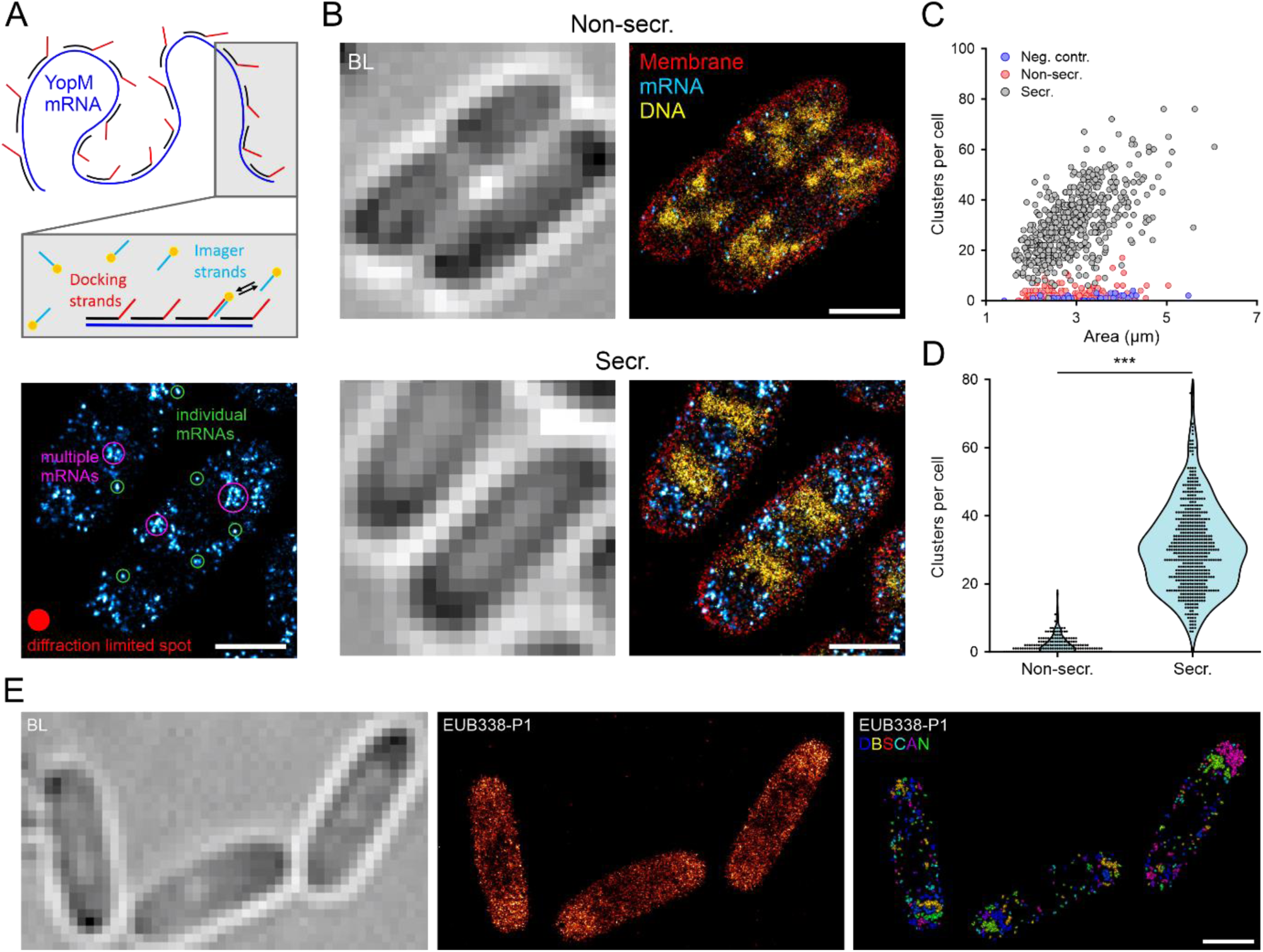
YopM mRNA localizes to the cytosol, but not to the membrane upon T3SS induction. **A)** Super-resolution single-molecule RNA fluorescence *in situ* hybridization (sr-smRNA-FISH) scheme (upper panel). The mRNA (*yopM*, blue) is targeted by short RNA-FISH oligos containing a non-hybridizing docking sequence (docking strand). RNA-FISH label positions are read out via DNA-PAINT using the transient hybridization of ATTO655-labeled imager strands. Resulting super-resolved images report on mRNA localization and clustering (lower panel). **B)** Representative multi-color super-resolution images of non-secreting and secreting cells. Membrane (red) and DNA (yellow) were visualized using PAINT, mRNA (cyan hot) using DNA-PAINT. **C)** Correlation between detected sr-smRNA-FISH clusters and cell size. *n*_replicates_ = 3, 4, 1 and *n*_cells_ = 176, 531, 33 for non-secreting, secreting cells and the negative control (no FISH probes), respectively. **D)** Quantification of sr-smRNA-FISH clusters in the indicated conditions (n as in C). Statistical analysis was performed using a Mann-Whitney-U test, ***, *p*<0.0001. **E)** sr-RNA-FISH of ribosomal RNA. Scale bars, 1 µm.

### T3SS activation alters chromosomal DNA organization

Upon activation of T3SS secretion, *Yersinia* cells display cell growth retardation [24,50] and cell elongation [51]. We confirmed the latter phenotype in our experimental setting by measuring single cell length, width and area in non-secreting and secreting cells, and determined that only the increased cell length contributes to the overall bigger cell area of secreting bacteria (**Fig. 4A-C**). Cell elongation and growth retardation could be explained by a defect in chromosome replication or segregation. To investigate this, we compared the DNA distribution in the previously tested conditions by DAPI staining. Secreting and non-secreting cells showed a drastically different DNA organization. The majority of non-secreting cells displayed two intensity spots at quarter positions, in line with a wild-type chromosomal segregation and comparable to what was observed in wild-type dividing cells at 28°C (**Fig. 4D**). In contrast, secreting cells showed the highest fluorescence intensity at mid-cell, independent of their cell length (**Fig. 4D**).

**Figure 4 –.**
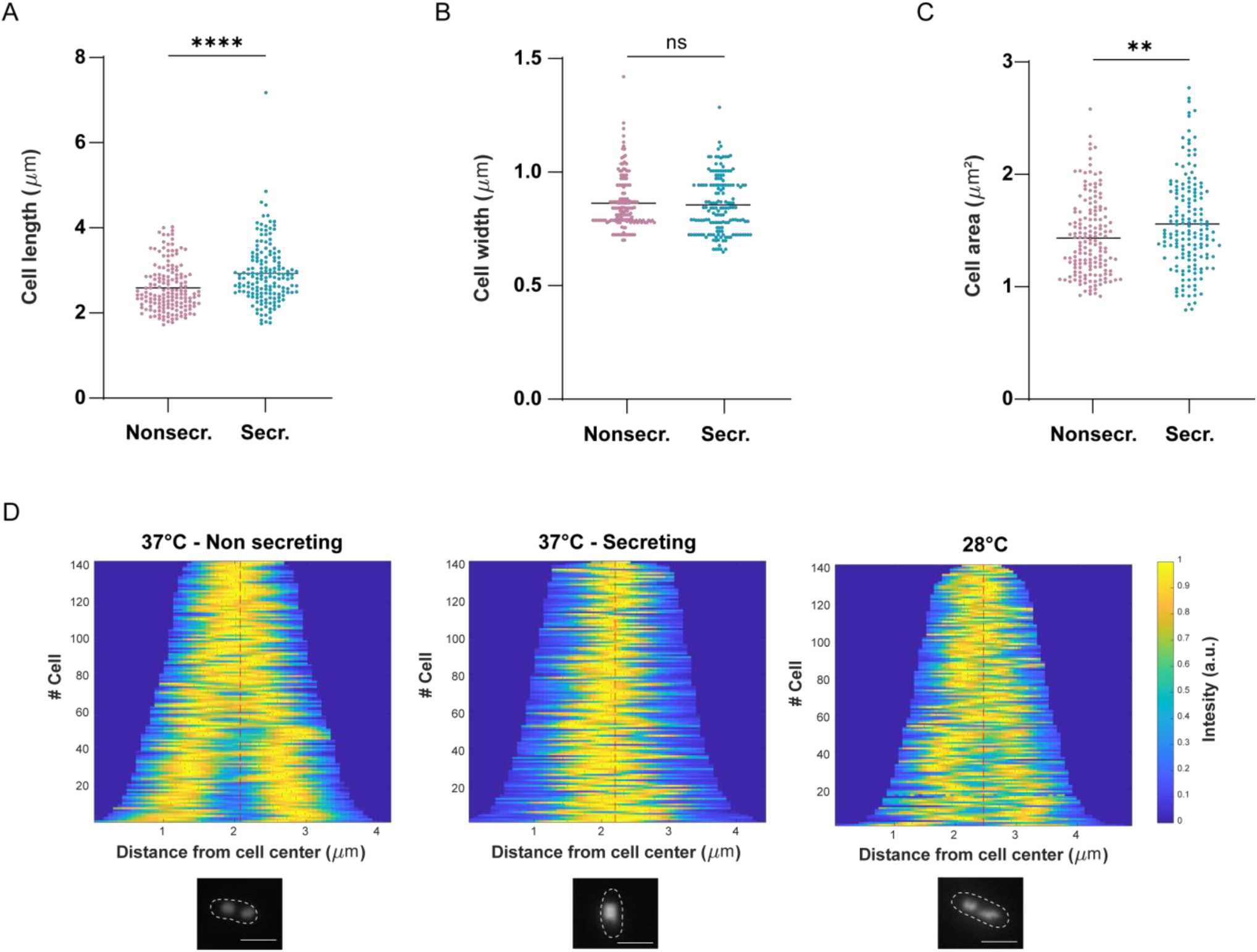
Cell length and DNA organization are affected by T3SS activation. **A) – C)** Cell length, width, and area measured in non-secreting (nonsecr., 37°C, Δ*sctD*) and secreting (secr., 37°C, Δ*sctW*) bacteria. Statistical analysis was performed using a non-parametric t-test, ****, *p*<0.0001; **, *p*<0.01 spots indicate single measurements. **D)** Chromosome organization of *Yersinia* wild-type cells at 37°C in non-secreting (Δ*sctD*) and secreting (Δ*sctW*) bacteria and 28°C in wild-type *Yersinia*. In demographs, cells are sorted according to cell length, DAPI signals are shown according to the intensity scale. Red spots represent intensity peaks. Lower panel, representative fluorescence microscopy images of cells stained with DAPI. Scale bar, 2μm. *n*=3 for all panels.

### Secretion leads to increased plasmid copy number and nucleoid rearrangement within 15 to 60 minutes

To temporally resolve the cellular rearrangements following T3SS activation, secretion was induced by Ca^2+^ chelation in a wild-type *Y. enterocolitica* strain, and pYV copy number and localization, cell length, and DNA organization were assessed as previously described (**Fig. 1B**, **2**, **4**) at the time of induction (T_0_) and 15 (T_15_), 30 (T_30_), or 60 min (T_60_) post-induction (**Fig. 5A**). pYV PCN gradually increased from the first time point tested, from an average value of 2.5 at T_0_ to a PCN of 4 at T_60_. As the full replication of the 70 kb pYV takes a few minutes (2 min at a replication speed of 600 bp/s), we assume that initiation of pYV replication is rapidly initiated upon activation of secretion.

**Figure 5 –.**
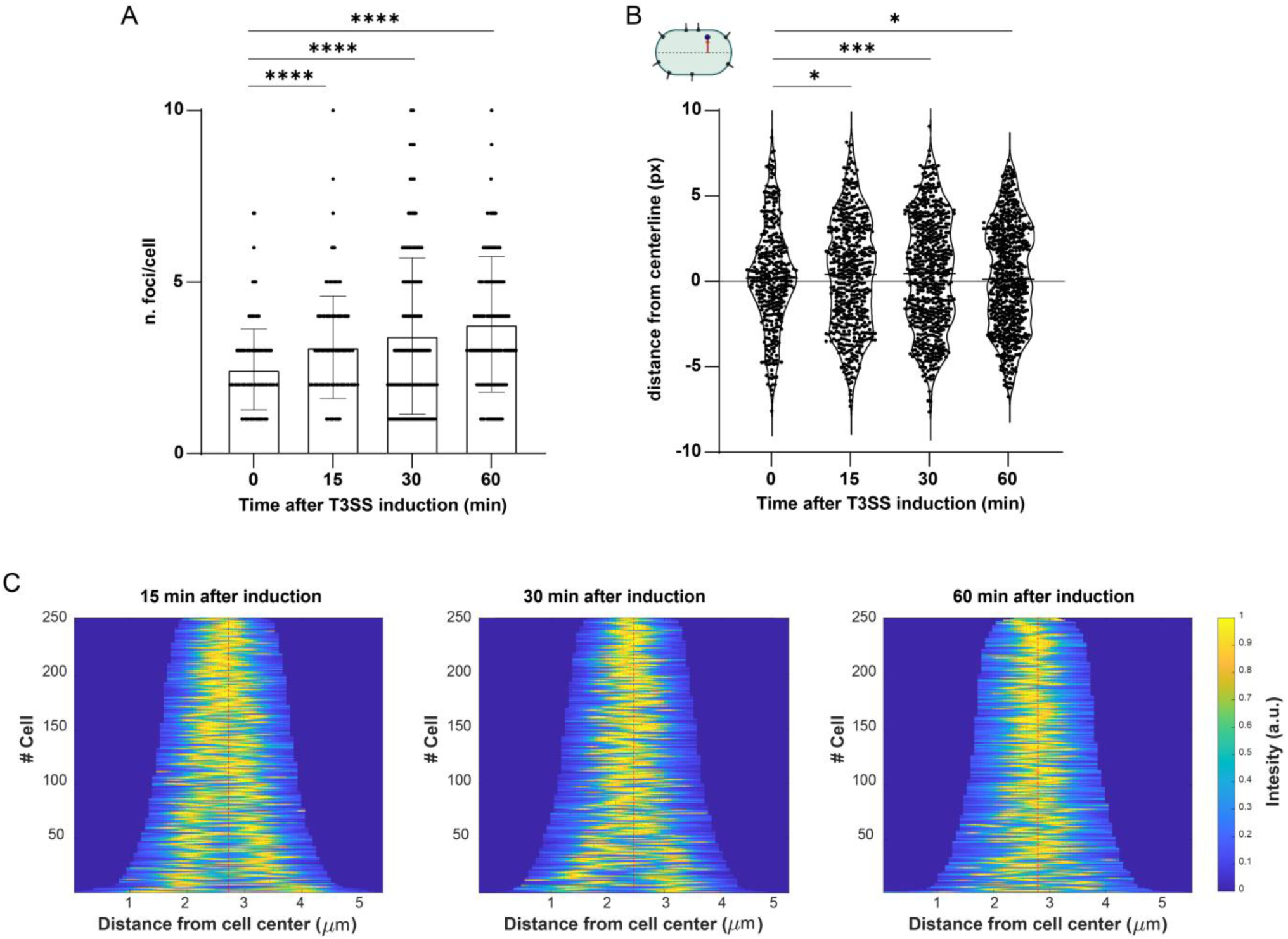
Kinetics of cellular rearrangement upon T3SS induction. **A)** Quantification of pYV (*yopE_1-53_-parS_P1_*) spots per bacterium, visualized by fluorescence microscopy before and 15, 30, or 60 min after T3SS induction by Ca^2+^ chelation at 37°C. Statistical analysis was performed using a non-parametric t-test, ****, *p*<0.0001. Each dot represents a single bacterium. **B)** Single-cell pYV plasmid localization (*yopE_1-53_-parS_P1_*) measured as distance from the centerline (px) in the conditions described in A. Statistical analysis was performed using a non-parametric t-test, *, *p*<0.05, ***, *p*<0.001. A) and B), each dot represents a single localization. **C)** Cell length and chromosome organization of *Yersinia* cells in the conditions described in A. In demographs, cells are sorted according to cell length, DAPI signals are shown according to the intensity scale. Red spots represent intensity peaks. *n*=3 for all panels.

Plasmid localization measured as the distance from the longitudinal centerline showed a more peripheral distribution at T_60_ than at T_0_. As for pYV PCN, plasmid localization was significantly shifted from the first time point tested (T_15_) compared to T_0_ (**Fig. 5B**). Importantly, a clear transition from a bipolar quarter-cell DNA distribution, typical of replicating and dividing cells, to a mid-cell DNA localization occurred upon the activation of secretion (**Fig. 5D**).

## Discussion

The T3SS is an essential virulence factor for a broad range of Gram-negative bacteria. To survive the innate immune response and to invade the host, pathogenic bacteria inject T3SS effector proteins directly from the bacterial cytoplasm into the target host cell. Regulation of the T3SS is essential due to the immunogenic effect of some components and the possible metabolic burden induced by machinery components and effectors synthesis. The link between T3SS secretion and bacterial physiology has always been of great interest, as the observation that virulent *Yersinia* cells display growth impairment (secretion-associated growth inhibition, SAGI) was pivotal to discovering the machinery [24,52]. Despite the striking phenotype, the nature of the link between T3SS secretion and bacterial growth is still highly debated [53–55].

To shed light onto this mechanism, we investigated key cellular parameters of secreting and non-secreting *Y. enterocolitica* at the single cell-level. The proposed role of SAGI as a strategy to preserve the energetic status of the cell is based on the observation that T3SS machinery components and especially the effectors are significantly upregulated upon secretion induction (reviewed in [10]). In *Y. pseudotuberculosis,* it was shown that during secretion, this upregulation correlates with an increase in pYV plasmid-copy number (PCN) [17,18]. We confirmed and further characterized this phenotype in *Y. enterocolitica* using a ParB/*parS*-based pYV labeling system (based on [56]) that allows to visualize and localize the plasmids as fluorescent spots (**Suppl. Fig. 1**). Single-cell detection showed a large heterogeneity of PCN in non-secreting and secreting bacteria, with secreting cells containing one to more than ten virulence plasmids (**Fig. 1**). As previously observed for *Y. pseudotuberculosis* [17], pYV PCN was increased by both temperature shift to 37°C (equivalent to host entry) and activation of the T3SS, leading to two sequential PCN upregulation steps (**Fig. 1**). Surprisingly, we found that virulence plasmids relocalize towards the membrane and the bacterial poles in secreting bacteria (**Fig. 2**). This phenotype is not a passive consequence of PCN increase as it was also observed the subset of T3SS-active cells that had the same pYV number as their non-secreting counterparts (**Fig. 2BC**). A possible reason for the relocation towards the membrane is transertion, the coupling of transcription, translation, and insertion of membrane proteins, which generates a dynamic link between the gene locus and the membrane [41]. A recent study showed initial evidence for transertion during T3SS-2 assembly in *Vibrio parahaemolyticus* [22]. The transcriptional activator of this T3SS, which unusually is a transmembrane protein, captures the T3SS gene locus and therefore leads to the transcription of T3SS components and effectors at the membrane [22]. Since in *Y. enterocolitica,* T3SS genes are encoded on the extra-chromosomal pYV plasmid, transertion would lead to a membrane localization of the pYV, which could occur either when the T3SS is assembled (shift to 37°), or upon activation of secretion, when large amounts of effectors are synthesized and exported [57]. As effectors have an exclusive role in the target cell and only need to be synthesized upon T3SS activation, they are prime candidates for transertion, particularly those which do not have a dedicated chaperone. We therefore investigated whether transertion is involved in the synthesis of YopM, a T3SS effector protein that belongs to this subgroup [43,44]. Investigating the subcellular localization of *yopM* transcripts using super-resolution microscopy revealed a strong upregulation under secreting conditions (**Fig. 3**). While the measured transcript abundance of 31 ± 14 copies/cell at 90 min after secretion activation is likely an underestimation due to technical reasons, the upregulation is striking. Co-imaging of DNA and bacterial membranes revealed that *yopM* mRNA is excluded from the nucleoid, but does not show a specific membrane enrichment (**Fig. 3A**). We thus conclude that transertion is not required for the secretion of this type of effector proteins.

An alternative explanation for the strong movement of pYV plasmids towards the membrane and bacterial poles is that increased T3SS gene expression during secretion acts as a driving force for the pYV plasmids and transcription-translation machinery from the nucleoid region [58,59]. It has been recently shown that enhanced extra-chromosomal gene expression induces ribosome accumulation at the synthesis site, leading to DNA compaction and defects in chromosome segregation [49]. Like *E. coli*, *Yersinia* does not harbor any known active ParAB*S* system to assist chromosome segregation, leaving the driving forces of this process open. To determine if the upregulation of pYV gene expression during secretion is accompanied by a relocalization of chromosomal DNA in *Yersinia*, we localized DNA in T3SS-active cells with high resolution using PAINT [47]. Live-cell imaging of DAPI-stained DNA showed a markedly altered DNA distribution upon secretion activation, changing from a largely bifocal one quarter/three quarter positioning to a unifocal central positioning, independently of the cell size (**Fig. 4D**).

To assess the dynamics of T3SS-induced cellular rearrangements, we investigated the temporal resolution of the previously discussed phenotypes. Interestingly, pYV plasmid copy number and localization significantly changed already after 15 min of T3SS induction, concomitantly with chromosome reorganization (**Fig. 5**). A fast response of pYV plasmid and consequent chromosome rearrangement would be compatible with reorganizations in the transcription/translation machinery. Large molecular assemblies, such as highly transcribed genes, were found to localize to the nucleoid surface to maximize entropy [60,61]. Upon secretion induction, genes localized on the pYV get highly transcribed, likely forming large molecular assemblies that segregate from the nucleoid region similar to highly transcribed genes from the nucleoid core. This is in agreement with a study by Martin *et al*. [62], in which relocation of *rrn* operons from the chromosome to a plasmid led to the formation of RNA polymerase assemblies at the cell poles. In addition to pYV relocalization, effector mRNAs are likely highly translated, as T3SS effectors account for a large fraction of the proteome under secreting conditions. Such mRNAs are thus expected to be localized in ribosome-dense regions of the cell [48]. This is in agreement with our super-resolution images, which show an enrichment of *yopM* mRNA and ribosomes at the polar cell regions. Consequently, the increased concentration of molecular crowders could then drive chromosome compaction, as demonstrated by Wu et al., who induced rapid nucleoid compaction by increasing molecular crowding in *E. coli* cell using an osmotic shock [63]. Central localization of chromosomal DNA prevents cell divison by nucleoid occlusion [64,65], which may be a causing or contributing factor for the SAGI [66,67].

Taken together, our data highlight the extent and nature of DNA rearrangement in secreting bacteria and provide a possible reason for the SAGI (**Fig. 6**), which – more than six decades after the discovery of the T3SS through this phenotype – is still a conundrum. Further single-cell studies of these events in other bacteria may help to discern the conservation of this striking phenotype and potential species-specific adaptations.

**Figure 6 –.**
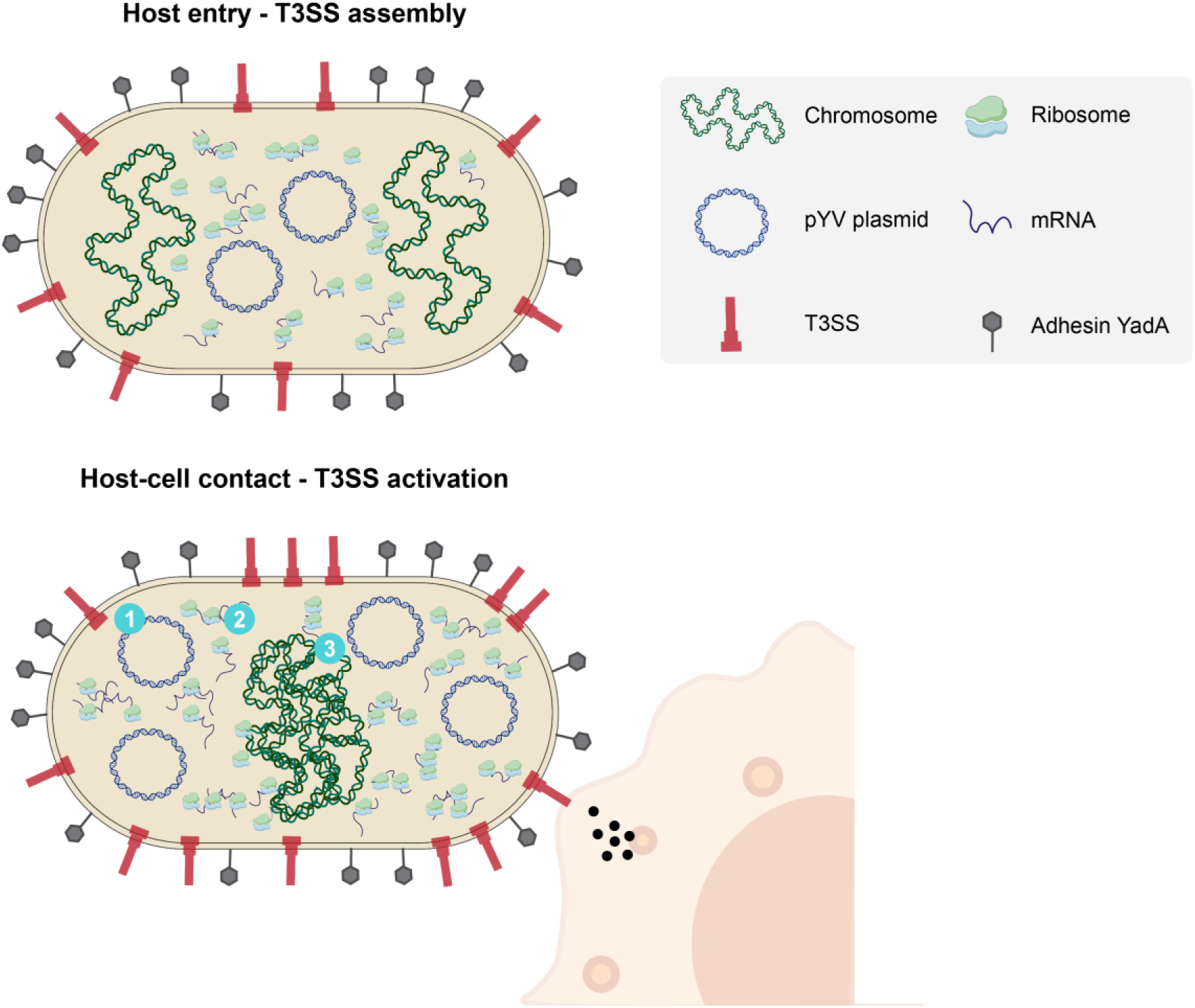
Influence of T3SS secretion on bacterial cell architecture. *Yersinia* relies on the adhesion protein YadA (depicted in grey on the bacterial surface) and effector secretion by the T3SS (depicted in red on the bacterial surface) to colonize the host by fighting the host immune system. Upon host entry, in the absence of host-cell contact (upper panel), the T3SS machinery is assembled, but secretion remains inactive [68]. At this stage, T3SS gene transcription is kept to a basal level, and pYV PCN (circular DNA depicted in blue) is on an average about 1-2 per number of chromosomal equivalents. Host-cell contact and T3SS activation (lower panel) lead to increased assembly of T3SS and strong upregulation of T3SS effector expression [11,69]. Simultaneously, the pYV PCN increases and the plasmids are excluded for the nucleoid, possibly due to the upregulation of T3SS genes (①). Under these conditions, effector transcripts (depicted as single filament in blue) accumulate in the polar regions, but without specific enrichment at the membrane. Increased expression of T3SS genes induces redistribution of the transcription and translation machinery (ribosomes depicted in light blue and green) away from the chromosome (depicted in green as a double helix string) (②), possibly preventing or reverting chromosome segregation (③) [49], which might then block cell division.

## Materials and methods

### Bacterial strain generation and genetic constructs

The *Y. enterocolitica* strains used in this study are based on the *Y. enterocolitica* wild-type strain MRS40. A list of strains and plasmids used in this study can be found in **Suppl. Tables 2-3**.

### Bacterial cultivation, secretion assay, and protein detection

*Y. enterocolitica* overnight cultures were grown on a shaking incubator at 28°C in brain heart infusion medium (BHI) supplemented with nalidixic acid (Nal, 35 µg/ml). Day cultures (BHI supplemented with 35 µg/ml Nal, 20 mM MgCl_2_, 0.4% glycerol, and 5 mM EGTA for secreting conditions or 5 mM CaCl_2_ for non-secreting conditions) were inoculated from stationary overnight cultures to an OD_600_ of 0.1 by directly adding the required amount of the overnight culture to fresh medium. To select for expression plasmid maintenance, 200 µg/ml ampicillin was added, where required. Day cultures were incubated at 28°C for 90 min after the inoculum. Expression of the *yop* regulon was then induced by a rapid temperature shift to 37°C. Where needed, protein expression from the pFHCP1-ParB_P1_-mScarlet plasmid was induced at the temperature shift by adding 100 µM Isopropyl β-D-1-thiogalactopyranoside (IPTG), as described for the respective experiments. Unless specified otherwise, bacteria were incubated for 150 min after the shift to 37°C. For secretion assays, 2 ml of the culture supernatant was harvested by centrifugation (10 min, 21,000 *g*) 180 min after the shift to 37°C. Proteins in the supernatant were precipitated with 10% trichloroacetic acid (TCA) at 4°C for 1-8 h. Precipitated proteins were collected by centrifugation (15 min, 21,000 *g*, 4°C). After washing with ice-cold acetone (1 ml), the pellets were resuspended in SDS-PAGE loading buffer and normalized to 0.3 OD units (ODu)/15 µl for total cell analysis, or 0.6 ODu/15 µl for secretion assays (1 ODu = 1 ml of culture at OD_600_ of 1, ~ 5 × 10^8^ *Y. enterocolitica*). After resuspension, samples were incubated for 5 min at 99°C and 15 µl were applied to SDS-PAGE analysis. Protein separation was performed on 15% SDS-PAGE gels and protein sizes were determined using the BlueClassic Prestained Marker (Jena Biosciences) as standard. For visualization, the gels were stained with FastGene-Q-stain (NipponGenetics). For immunoblots, the separated proteins were transferred to a nitrocellulose membrane. Primary rabbit anti-mCherry antibody (BioVision 5993, 1:2,000) was used in combination with a secondary anti-rabbit antibody conjugated to horseradish peroxidase (Sigma-Aldrich A8275, 1:10,000). For visualization, Immobilon Forte chemiluminescence substrate (Sigma-Aldrich) was used in a LAS-4000 Luminescence Image Analyzer.

### Fluorescence Microscopy

For fluorescence microscopy, bacteria were grown as described above. After 2.5 h at 37°C, 500 µl of culture were harvested by centrifugation (2 min, 2400 *g*) and resuspended in 250 µl of microscopy medium (100 mM 2-[4-(2-Hydroxyethyl)piperazin-1-yl]ethane-1-sulfonic acid (HEPES) pH 7.2, 5 mM (NH_4_)_2_SO_4_, 100 mM NaCl, 20 mM sodium glutamate, 10 mM MgCl_2_, 5 mM K_2_SO_4_, 50 mM 2-(N-morpholino) ethane sulfonic acid (MES), 50 mM glycine). 2 µl of bacterial resuspension were spotted on an agarose pad (1.5% low melting agarose (Sigma-Aldrich) in microscopy medium, 1% casamino acids, 5 mM EGTA) in a glass depression slide (Marienfeld). For imaging, a Deltavision Elite Optical Sectioning Microscope equipped with a UPlanSApo 100×/1.40 oil objective (Olympus) and an EDGE sCMOS_5.5 camera (Photometrics) was used. z stacks with 9 slices (Δz = 0.15 µm) were acquired. The mScarlet signal was visualized using a mCherry filter set (excitation: 575/25 nm, emission: 625/45 nm) with 0.2 s exposure time. Images were processed with FIJI (ImageJ 1.51f/1.52i/1.52n) [70]. Fluorescence quantification was performed in FIJI. For visualization purposes, selected fields of view adjusted identically for brightness and contrast within the compared image sets are shown.

For DNA staining, after 2.5 h at 37°C, 1 ml of cells were harvested and resuspended in 50 µl fresh minimal medium supplemented with 1 μg/ml DAPI. After 3 min incubation at 37°C in the dark, cells were pelleted and resuspended in 100 µl and imaged as previously described. The DAPI signal was visualized using the DAPI filter set (excitation: 390/18 nm, emission: 435/48 nm) with 0.2 s exposure time. For data analysis and demograph generation, individual *Y. enterocolitica* cells were first segmented using the neural network StarDist [71]. Segmented cells were loaded into BacStalk [72] for cell length, width, area, and DAPI signal quantification. Single-cell maximal DNA intensity was represented according to cell length.

### Quantification and localization of fluorescent pYV foci

To quantify the number of pYV spots, a mScarlet-I fused ParB_P1_ (mScarlet-I-ParB_P1_) cloned under an IPTG-inducible promoter was expressed from plasmid in a strain harboring the correspondent *parSP1* sequence in the pYV plasmid (*YopE_1-53_-parS_P1_*). For fluorescent spot visualization, *Y. enterocolitica* cells were first segmented using the neural network StarDist [71]. Spot detection and localization were performed on segmented cells in Oufti [73] with the following spot detection parameters optimized for the dataset. Data were then analyzed in Matlab (MATLAB R2020a) and the spot localization was displayed as distance from the centerline (d in pixels, pixel size = 64.8 µm^2^). To determine the localization of foci along the longitudinal axis, fluorescent foci, the cell center, and cell poles were manually classified in a set of images for which the conditions were blinded, and the relative location was determined as the ratio of the distances of cell center to spot and cell center to respective cell pole.

### mRNA Fluorescence *in situ* hybridization

Cells were grown for mRNA Fluorescence in situ hybridization (FISH), as previously described. After 2.5 h at 37°C, cells were prepared according to [74]. 3 × 10^9^ cells were fixed directly by adding formaldehyde (3.7 % v/w) and incubated for 30 min, gently mixing. The cell pellet was washed twice with 1 ml PBS (3.5 min, 600 *g*) and then resuspended for permeabilization in 70% Ethanol and incubated while mixing gently. After 1 h, the cell pellet was resuspended in 1 ml washing solution (formamide 40% v/w, saline-sodium citrate (SSC) buffer 2x) and mixed for 5 min. Cells were pelleted (7 min, 600 *g*) and resuspended in 50 µL hybridization solution (in 10 ml, 1g dextran, 3.530 ul formamide, 10 mg *E.coli* tRNA, 1 ml 20x SSC, 200 mM vanadyl ribonucleoside complex (VRC) and 50 mg/ml BSA) mixed with the probe solution (1 µM). Samples were incubated O/N at 30°C. After O/N hybridization with the probes, 10 µl of the sample was washed two times with 200 µl of wash solution, with a 30 min incubation at 30°C with the solution before the second centrifugation. The washing steps are repeated two times to ensure the removal of any probe traces. For sample immobilization and preparation for imaging 10 µl of the hybridization mixture were washed thrice with washing buffer (2x SSC, 40% formamide) for 30 min at 30°C. All steps were performed with RNase-free solutions and in RNase-free environment. Cells were finally resuspended in 150 µl RNase-free PBS and immobilized on KOH-cleaned (3 M, 1 h) and PLL-coated chamber slides. Chambers were washed twice with RNase-free PBS to remove non-adherent cells. Fiducial markers (90 nm gold nano-urchins,Cytodiagnostics) were added and allowed to settle for 30 min, before washing and blocking the cells with 2% RNase-free BSA (Carl Roth 3737.2) for 30 min. Finally, cells were washed with RNase-free PBS twice and stored at 4°C or imaged.

All single-molecule imaging experiments were performed on a commercial N-STORM system (Nikon Instruments) equipped with a 100x Apo TIRF oil objective (NA 1.49) and an Andor iXon Ultra 897 EMCCD camera. For DNA-PAINT imaging, 0.5-5 nM P1-ATTO655 imager strand in imaging buffer (100 mM Tris pH 8.0 + 500 mM NaCl) was added to the samples. Samples were imaged in HILO mode using 1 kW/cm^2^ 640 nm illumination, an exposure time of 150 ms, a readout rate of 1 MHz, a preamplifier of 1 and an EM gain of 50. 30,000 frames were acquired for FISH-PAINT imaging. Conventional PAINT imaging of DNA and membrane was performed using JF_646_-Hoechst (200 – 400 pM) and Nile Red (200 – 400 pM) in DNA-PAINT imaging buffer to maintain osmolality and prevent cell shrinkage. In contrast to DNA-PAINT imaging, the EM-gain was increased to 200-400, using a readout rate of 10 MHz.

SMLM images were analyzed using Picasso [75]. Drift correction was performed using RCC with a window size of 500 – 1000 frames. Subsequent frame localizations were linked using a radius of 1 px and allowing for 1 dark frame. PSF width σ in x and y was filtered according to 64 nm < σ_x/y_ < 208 nm (Nile Red) or 80 nm < σ_x/y_ < 224 nm (ATTO655 and JF_646_-Hoechst). Images were registered in Fiji using cell boundaries extracted from the Nile Red image.

### Counting of mRNA clusters

The number of mRNA clusters in individual cells were determined using the DBSCAN algorithm implemented in the Picasso software. DBSCAN analysis requires two parameters, namely the search radius ε and the minimum number of molecules in the search radius required for the localization to be attributed to a cluster (minpts). We set ε to twice the value of the experimental localization precision (determined via NeNA analysis, 10.6 nm). minpts was determined by manually picking 200 – 300 apparent mRNA clusters (see Fig. 3A, green circles) and calculating the median value of localizations contained in each picked region. Of note, we performed this analysis on localization files not corrected for subsequent-frame localizations (linking). The reason for this is that true DNA-PAINT signals have a longer binding time than the unspecific binding of imager strands, leading to multiple localizations and an improved signal-to-background ratio in the reconstructed image. To remove clusters that origin from single, very long binding events (which result in cluster with many localizations), we filtered the DBSCAN-derived clusters by the standard deviation of the frame number (frame s.d.). If all localizations occur in a small time window, the frame s.d. is low, while recurring binding events over the entire acquisition period lead to a high frame s.d.. We determined a cutoff value of 5,000 frames for the frame s.d. based on the negative control (secreting conditions, no FISH probes added during sample preparation). Finally, we rendered an image in which each true cluster adds 1 gray value to the pixel coordinates of its centroid. Finally, the number of clusters were determined by measuring the intensity within the cell outlines. Key steps of the pipeline are shown in **Suppl. Fig. 5**. Statistical analysis of cluster numbers and densities was performed with Origin 2023b (OriginLab).

### qPCR

Total genomic DNA of *Y. enterocolitica* cells grown as described above (2 × 10^9^ cells) was extracted using the PureLink Miniprep Kit (Thermofisher Scientific) according to the manufactureŕs protocol. qPCR reactions were performed on an Applied Biosystems 7500 Real-Time PCR system using the Luna Universal qPCR MasterMix (New England Biolabs) and the primers listed in **Suppl. Table 1**. Data analysis was performed using the comparative threshold cycle (C_T_) method [76].

## Notes

### Competing Interest Statement

The authors have declared no competing interest.

