## Supporting Information for "Activation of the *Yersinia* type III secretion system induces large-scale chromosomal and virulence plasmid DNA rearrangements"

for

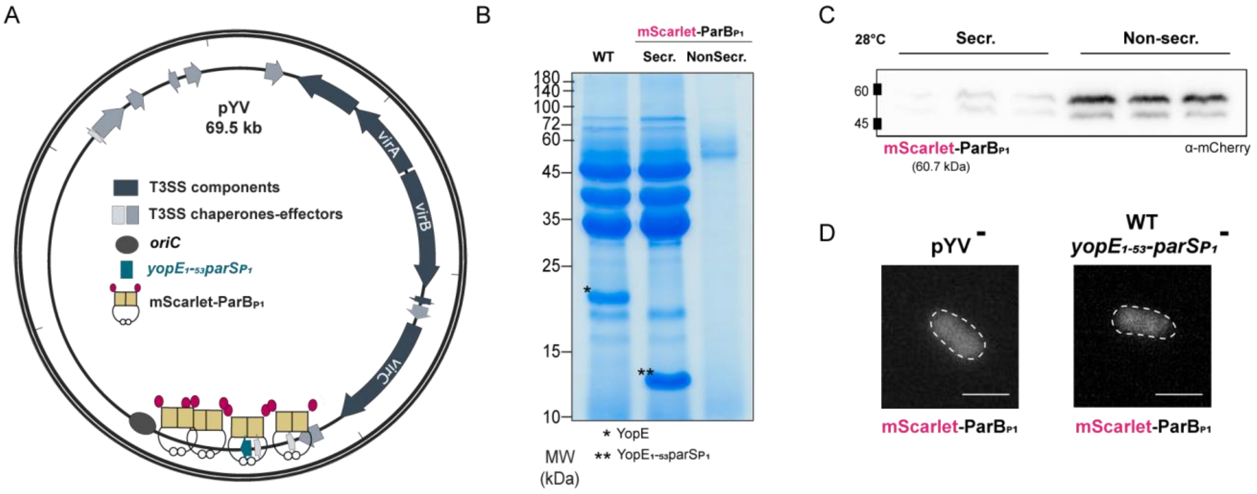


**Supplementary Figure 1 – Establishment of ParAB*S* P1 labeling system in *Y. enterocolitica* for pYV labeling**

**A)** Schematic display of the genes encoding for T3SS structural components (operons *virA*, *virB*, *virC*, and translocator operon (transl.), dark grey), T3SS effectors and chaperones (lighter shades of grey), *oriC* (grey circle) and $yopE_{1\text{-}53}$-$parS_{P1}$ (pink) on the pYV virulence plasmid. To exclude any effect on the native replication of the pYV plasmid, we introduced a heterologous ParBS system (derived from [1]) and expressed ParB from a plasmid. mScarlet-ParB$_{P1}$ binding to $yopE_{1\text{-}53}$-$parS_{P1}$ and sliding is represented. **B)** Secretion assay displaying the effectors secreted by the T3SS in wild-type (WT), ParABS P1 labeled secreting (secr.) and non-secreting (non-secr.) strain background. mScarlet-ParB$_{P1}$ was expressed from plasmid (100 µM IPTG). **C)** Expression of mScarlet-ParB$_{P1}$ (expected molecular weight 60.7 kDa) expressed from plasmid (100 µM IPTG) in *Y. enterocolitica* at 28°C in secreting (secr.) and non-secreting (non secr.) background. Immunoblot using antibodies directed against mCherry as indicated. **D)** Sample micrographs of fluorescence microscopy detecting mScarlet-ParB$_{P1}$ signal in the indicated genetic backgrounds; scale bar, 2 µm. *n*=3 for all panels.

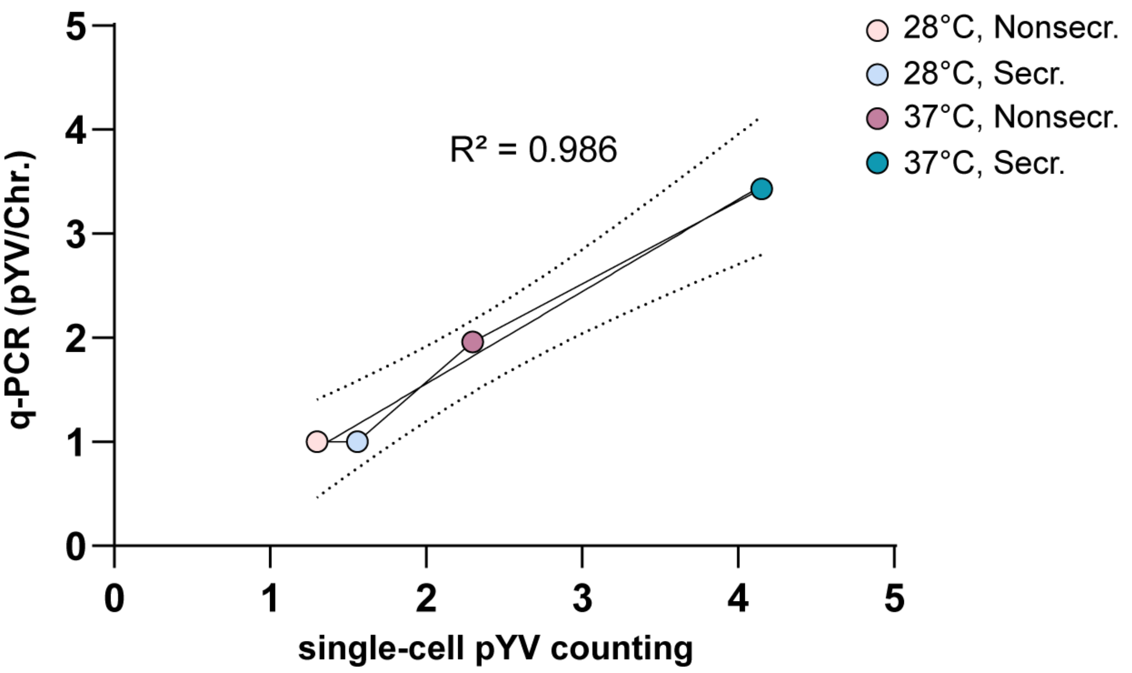


**Supplementary Figure 2 – Cross-validation of pYV quantification strategies**

Linear regression analysis of pYV quantification data obtained through quantitative PCR (pYV/chromosomes, more precisely *virF*/*gyrB*), and single pYV foci counting using the ParB-*parS*$_{P1}$ labeling system. The single spots represent the interpolations of the quantifications obtained from the two techniques at the conditions tested: non-secreting (Δ*sctD*) and secreting (Δ*sctW*) at 28°C and 37°C. Light pink, non-secreting 28°C; light blue, secreting 28°C; dark pink, non-secreting 37°C; dark blue, secreting 37°C. The coefficient of determination ($R^2$) is displayed.

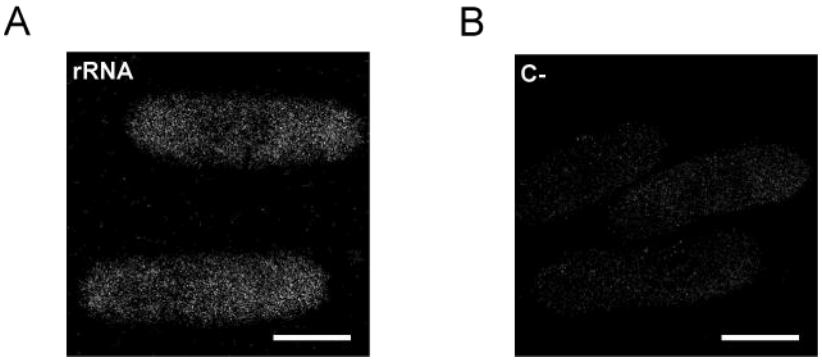


**Supplementary Figure 3 – Controls for RNA-FISH experiments**

Representative images of controls for RNA-FISH imaging pipeline, using **A)** a eubacteria-specific rRNA probe (EUB338) as a positive control (rRNA, see Fig. 3E) and **B)** a sample processed without any mRNA targeting probe as a negative control (C-).

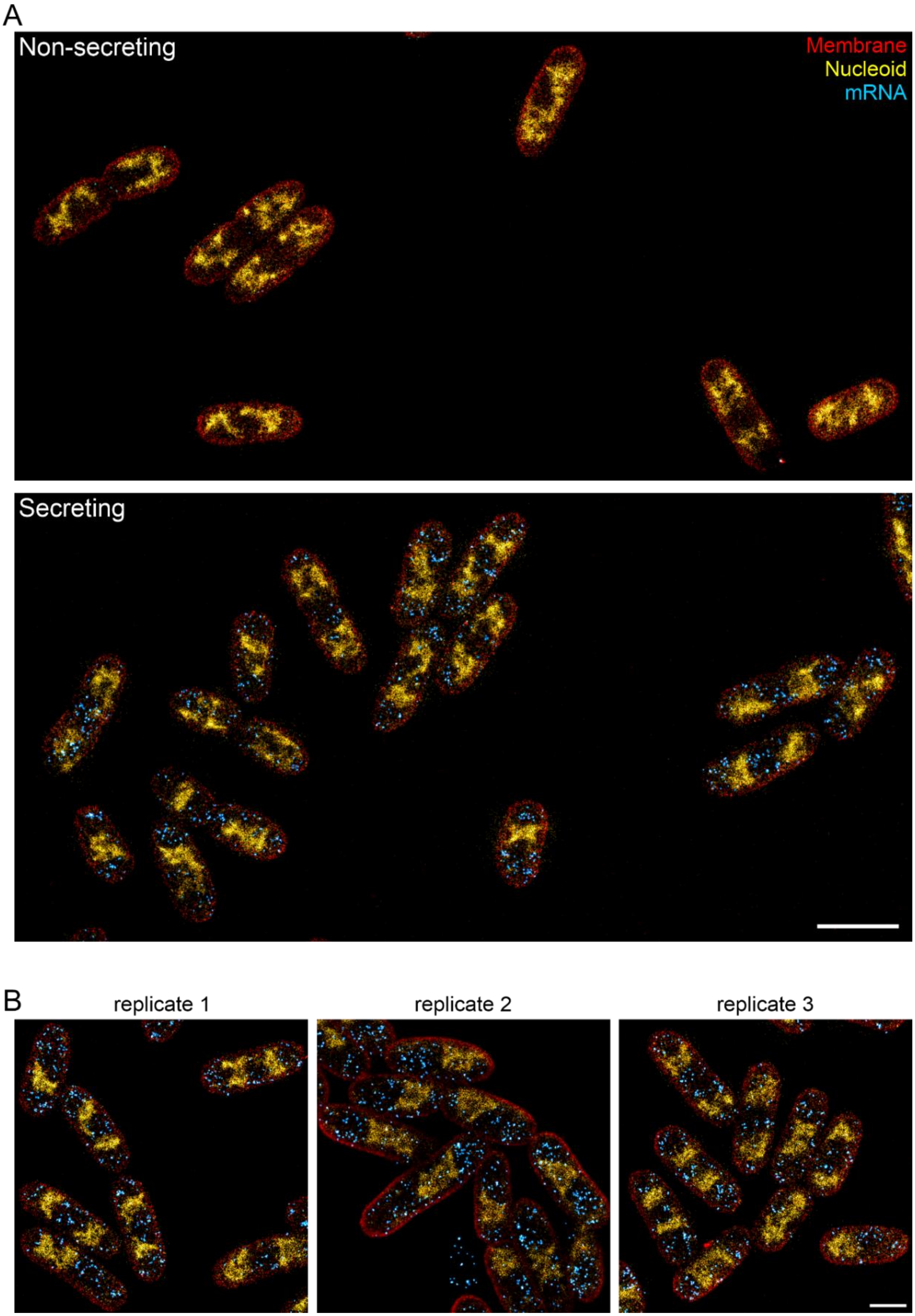
A
Non-secreting
Membrane
Nucleoid
mRNA
Secreting
B
replicate 1
replicate 2
replicate 3

*previous page:* **Supplementary Figure 4 - Multi-color imaging of *Y. enterocolitica***

Membranes (red) were imaged using Potomac Gold, DNA using $JF_{646}$-Hoechst and mRNA using DNA-PAINT. **A)** Larger fields of views for non-secreting and secreting conditions. **B)** Representative regions of different replicates under secreting conditions. Scale bars are 2 µm (A) and 1 µm (B).

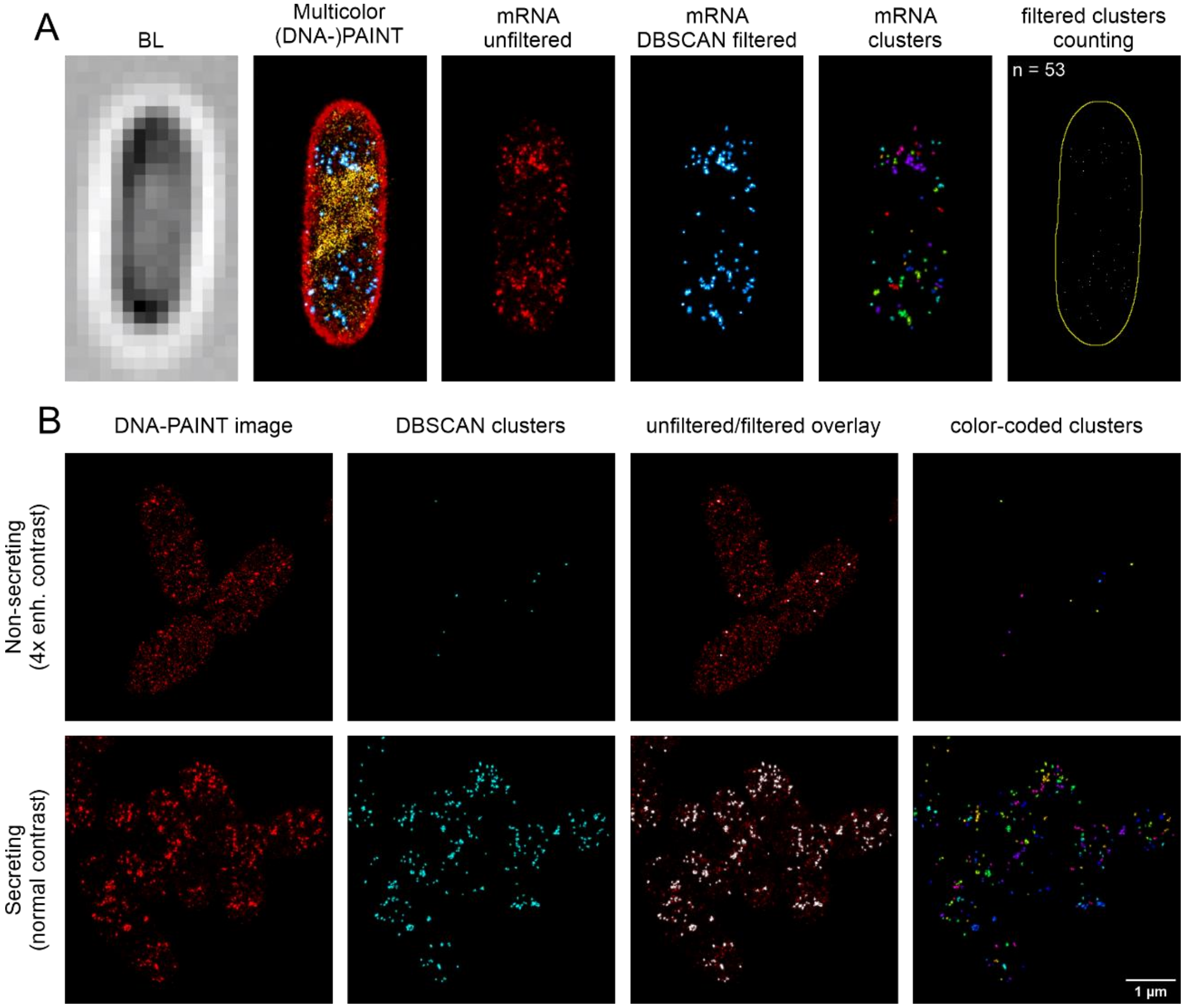


**Supplementary Figure 5 - Analysis pipeline for the quantification of sr-smRNA-FISH clusters**

**A)** Procedure shown for an individual cell grown under secreting conditions. The DNA-PAINT image (3rd image, mRNA unfiltered) is analyzed for localization clusters using DBSCAN, resulting in the removal of background or small (artifactual) clusters (4th image). A color-rendered image of the detected clusters is shown in the 5th image. Clusters are then filtered by kinetic properties and an images is rendered with cluster centers adding 1 gray value to the position of their centroid (6th image). Clusters are then counted by measuring the total intensity within cell outlines (yellow line) that were determined in the super-resolved membrane image or brightlight image. **B)** Exemplary DBSCAN cluster analysis for non-secreting and secreting cells. Non-secreting cells are displayed with 4x increased contrast for better visibility.

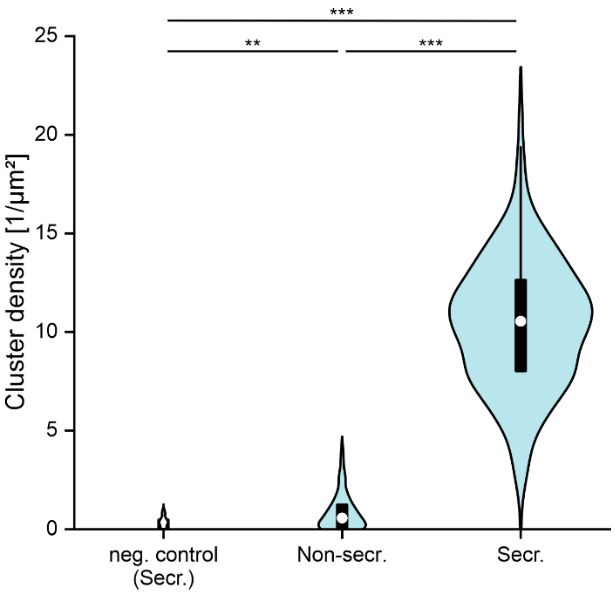


**Supplementary Figure 6 - Distribution of cluster densities of individual cells under different conditions**

The negative control represents cells grown under secreting conditions, which were prepared without addition of RNA-FISH probes, but treated and images similarly otherwise. Statistical analysis was performed using a Mann-Whitney-U test, **, p<0.001, ***, p<0.0001. n = 33, 176 and 531 cells.

### Suppl. Table 1 - Primers used in this study

| Primer name | Sequence (5´→ 3´) | Used for |
|---|---|---|
| AD1489 | GACTAGATCTCAGGATGCCGAAGAGCATCC | pFE020 |
| AD1490 | GACTGAATTCTCCGCGAAATTATGAGTCACGAA | pFE020 |
| AD1977 | TCACCAACAACATTCCACAG | qPCR *gyrB* |
| AD1978 | TTCGACCGCAGTTTTTACC | qPCR *gyrB* |
| AD1983 | AGACGAGACAATGCCACAC | qPCR *virF* |
| AD1984 | GCAGAGCCGAGAGGAATAAAG | qPCR *virF* |

### Suppl. Table 2 - Plasmids used in this study

| Plasmids | Genotype | Reference |
|---|---|---|
| pKNG101 | *oriR6K sacBR*$^+$ *oriTRK2 strAB*$^+$<br>(suicide vector for homologous recombination) | [2] |
| pFE020 | pKNG101-YopE$_{1-138}$-ParS$_{P1}$-FLAG<br>(mutator for insertion of ParSP1-FLAG into *yopE* locus) | This work |
| pFHCP1-ParBP1-mScarlet | pFHCP1::ParBP1-mScarlet<br>(vector for ParBP1-mScarlet overexpression) | [1] |

### Suppl. Table 3 - Strains used in this study

| Strain | Genotype | Reference |
|---|---|---|
| MRS40 | Wild-type *Y. enterocolitica* E40 *ΔblaA* | [3] |
| IML421*asd* (*ΔHOPEMTasd*) | MRS40 *yopO*$_{\Delta 12-427}$ *yopE*$_{21}$ *yopH*$_{\Delta 11-352}$ *yopM*$_{23}$ *yopP*$_{23}$ *yopT*$_{135}$ *Δasd* | [4] |
| pIM41 | MRS40 *ΔsctW* | [5,6] |
| AD4051 | MRS40 *ΔsctD* | [6] |
| FE016 | MRS40 *yopE*$_{1-53}$*::parSP1* | This study |
| FE017 | MRS40 *egfp-sctQ yopE*$_{1-53}$*::parSP1 ΔsctW* | This study |
| FE018 | MRS40 *egfp-sctQ yopE*$_{1-53}$*::parSP1 ΔsctD* | This study |
| pYV$^-$ | MRS40 cured of virulence plasmid | [7] |
